# Evolution and clinical impact of genetic epistasis within EGFR-mutant lung cancers

**DOI:** 10.1101/117291

**Authors:** Collin M. Blakely, Thomas B.K. Watkins, Wei Wu, Beatrice Gini, Jacob J. Chabon, Caroline E. McCoach, Nicholas McGranahan, Gareth A. Wilson, Nicolai J. Birkbak, Victor R. Olivas, Julia Rotow, Ashley Maynard, Victoria Wang, Matthew A. Gubens, Kimberly C. Banks, Richard B. Lanman, Aleah F. Caulin, John St. John, Anibal R. Cordero, Petros Giannikopoulos, Philip C. Mack, David R. Gandara, Hatim Husain, Robert C. Doebele, Jonathan W. Riess, Maximilian Diehn, Charles Swanton, Trever G. Bivona

**Author notes:** These authors contributed equally to this work (Collin M. Blakely, Thomas B.K. Watkins, Wei Wu, Beatrice Gini). Corresponding authors: Trever G. Bivona MD PhD and Charles Swanton MD PhD.

## Introductory paragraph

The current understanding of tumorigenesis is largely centered on a monogenic driver oncogene model. This paradigm is incompatible with the prevailing clinical experience in most solid malignancies: monotherapy with a drug directed against an individual oncogenic driver typically results in incomplete clinical responses and eventual tumor progression^1-7^. By profiling the somatic genetic alterations present in over 2,000 cases of lung cancer, the leading cause of cancer mortality worldwide^8,9^, we show that combinations of functional genetic alterations, i.e. *genetic collectives* dominate the landscape of advanced-stage disease. We highlight this polygenic landscape and evolution of advanced-stage non-small cell lung cancer (NSCLC) through the spatial-temporal genomic profiling of 7 distinct tumor biopsy specimens and 6 plasma specimens obtained from an *EGFR*-mutant NSCLC patient at (1) initial diagnosis of early-stage disease, (2) metastatic progression, (3) sequential treatment and resistance to 2 EGFR inhibitors, (4) death. The comprehensive genomic analysis of this case, coupled with circulating free (cf) tumor DNA profiling of additional advanced-stage EGFR-mutant NSCLC clinical cohorts with associated treatment responses uncovered features of evolutionary selection for multiple concurrent gene alterations: including the presence of EGFR inhibitor-sensitive (*EGFR^L858R^;EGFR^exon19del^*) or inhibitor-resistant (*EGFR^T790M^;EGFR^C797S^*) forms of oncogenic *EGFR* along with cell cycle gene alterations (e.g. in *CDK4/6, CCNE1, RB1)* and activating alterations in *WNT/*β-catenin and *PI3K* pathway genes, which our data suggest can cooperatively impart non-redundant functions to limit EGFR targeted therapy response and/or promote tumor progression. Moreover, evidence of an unanticipated parallel evolution of both *EGFR^T790M^* and two distinct forms of oncogenic *PIK3CA* was observed. Our study provides a large-scale clinical and genetic dataset of advanced-stage EGFR-mutant NSCLC, a rationale for specific polytherapy strategies such as EGFR and CDK4/6 inhibitor co-treatment to potentially enhance clinical outcomes, and prompts a re-evaluation of the prevailing paradigm of monogenic-based molecular stratification for targeted therapy. Instead, our findings highlight an alternative model of *genetic collectives* that operate through epistasis to drive lung cancer progression and therapy resistance.

## Main text

The identification of specific, therapeutically targetable genetic alterations that drive the growth of many cancers including non-small cell lung cancer (NSCLC), the major subtype of lung cancer, has revolutionized the diagnosis and treatment of patients. The clinical success of genetically-targeted therapies such as EGFR, ALK, ROS-1 and BRAF inhibitors^1-7,1-10,11^ has led to the current monogenic driver oncogene model of tumorigenesis, wherein advanced-stage NSCLC patients are selected for a gene-targeted monotherapy based on the identification of an individual specific driver alteration present in the tumor (e.g. mutant *EGFR* or an *ALK* gene rearrangement). This model is incompatible with the prevailing clinical experience in most advanced-stage solid malignancies including NSCLC, wherein monotherapy with a drug directed against a particular oncogenic driver protein, while typically more effective than cytotoxic chemotherapy, generally results in incomplete responses and eventual tumor progression in patients (reviewed in^12^). In contrast to this monogenic model of disease pathogenesis and treatment, it is possible that advanced-stage NSCLCs exhibit a polygenic landscape of functional gene variants that cooperate to limit response to treatment and/or promote tumor progression, potentially in conjunction with other events (e.g. secreted factors and/or stromal cells). In support of this hypothesis, recent important studies have revealed intra-tumor heterogeneity and co-occurrence of gene alterations in small NSCLC cohorts^13-17^. However, these prior reports were limited by either the small clinical cohort size and lack of associated clinical and treatment outcome data, the number of genes analyzed, and/or were focused on early-stage (e.g. TCGA lung adenocarcinoma project) rather than advanced-stage NSCLC, the latter comprising the patient population which is treated with systemic targeted therapy and therefore of paramount relevance. On this basis, we set out to address a critical unresolved question in the field: to what extent are advanced-stage NSCLCs comprised of combinations of functional somatic genetic alterations (*genetic collectives*) that could operate through genetic epistasis to cooperatively influence tumor progression and clinical outcomes? Critical to addressing this question was our ability: (1) to assemble a substantial cohort of clinical samples linked to clinical outcome and treatment response data, many through longitudinal specimen collection in individual patients, and (2) to characterize the somatic genetic alterations present within these cancers by whole exome sequencing and/or cfDNA profiling and link the genetic findings to clinical outcomes.

To test the hypothesis that a polygenic landscape of functional gene variants can cooperate to limit response to treatment and/or promote tumor progression through genetic epistasis in advanced-stage lung cancer, we first obtained a unique series of *EGFR*-mutant NSCLC clinical specimens from an individual patient and profiled them by both tumor-based whole exome sequencing and cfDNA analysis. EGFR-mutant NSCLC represents a specific NSCLC subtype (typically comprising 10-20% of cases) that is defined by the presence of an oncogenic mutation in *EGFR* that identifies patients who are likely to respond to an approved EGFR inhibitor (erlotinib, gefitinib, afatinib, osimertinib). These specimens were obtained over the course of 6 years from a patient who was initially diagnosed with surgically resectable disease in 2009, then suffered local progression in the mediastinum and distant metastatic progression in the lungs, bone, and brain over the ensuing 6 years. This disease course included an incomplete response and then progression first upon treatment with the first-generation EGFR inhibitor erlotinib and then upon treatment with the third-generation EGFR inhibitor rociletinib^3^ (Fig. S1). Seven tumor (4 lung, 2 bone, and 1 lymph node) and six plasma specimens were analyzed longitudinally during this disease progression to identify somatic genetic alterations, including 4 specimens obtained at autopsy upon lethal tumor progression on rociletinib.

The whole exome sequencing analysis of the tumor samples showed multiple concurrent functionally-relevant somatic alterations present at the initial diagnosis of early-stage disease (R1), including clonal and truncal *EGFR^exon19del^, CTNNB1*^S37F 18^, *SMAD4^L146X^,* and *RBM10^S167X^* mutations and *CDK2NA* copy number loss (Fig. 1a-b, S2, S3, Tables S1-S2). Progression to mediastinal lymph node metastasis (R2) was marked by acquisition of mutations in *PRKCA*(N468I)^19^ and *PIK3CA*(G106V)^20^, in addition to copy number gain (CNG) in the genomic region encoding *EGFR, CDK6, MET,* and *BRAF* (Fig. 1a-c, S2, Tables S1, S3). Progression on erlotinib therapy occurred upon acquisition of the *EGFR^T790M^* mutation, as found in ~60% of EGFR-mutant NSCLC patients who progress on first-generation EGFR inhibitor treatment^21^, and the persistent presence of additional genetic events including *CTNNB1^S37F^* and *PIK3CA^G106V^.* Our data suggest that the *PIK3CA^G106V^* mutation arose prior to EGFR inhibitor treatment and also prior to acquisition of the *EGFR^T790M^* mutation (Fig. 1a). Interestingly, the *EGFR^T790M^* mutation may have arisen in this case through a previously unreported instance of independent dual clonal origins. Given that the *EGFR^T790M^* mutation was found in metastatic sites that harbored *PIK3CA^G106V^* (R3-left lung at erlotinib progression, R4-left lung at rociletinib progression, and R6-right lung at rociletinib progression) and those that did not (R5-right rib metastasis and R7-spine metastasis), the data suggest that the *EGFR^T790M^* mutation arose in parallel sites independently (Fig. 1a-c, S3). Further progression on rociletinib was associated with the emergence of additional subclonal genetic co-alterations, including *PIK3CA^H1047R^* (R5-right rib), *RB1^R857H^* (R4-left lung), *CHD4^H1151P^* (R6-right lung) and *TLR4^R289Q^* (R5-right rib) (Fig. 1a-c). The activating *PIK3CA^G106V^* mutation^20^ was not found in all of the post-rociletinib metastatic sites (present in R4, R6; absent in R5, R7), demonstrating lesion-specific metastatic heterogeneity (Fig. 1a-b). Intriguingly, a subclonal *PIK3CA*^H1047R 22^ oncogenic mutation was found in R5 (right rib, post-rociletinib), suggesting another parallel evolution of two different PIK3CA oncogenic mutations in this patient’s cancer (Fig. 1a-c, S3). Although an *RB1* somatic mutation (R847H) was detected in R4 (left lung-at rociletinib progression) and *RB1* inactivation has previously been associated with a transition from lung adenocarcinoma to small cell carcinoma in the setting of EGFR inhibitor resistance^23,24^, there was no evidence of histological transformation to small cell carcinoma in these clinical specimens perhaps due to the lack of a somatic *TP53* genetic alteration in this case (Fig. S4). While plasma samples for cfDNA analysis were not available for all initial clinical events (i.e. diagnosis and metastatic progression prior to erlotinib treatment), the coupling of the tumor biopsy and cfDNA exome data (Methods) in this patient revealed examples of ubiquitous (e.g. *EGFR^ex19del^, CTNNB1^S37F^*) and more lesion-restricted (*PIK3CA^G106V^, RB1^R857H^, and TLR4^R289Q^*) mutations in the plasma (Fig. 1d). This finding suggests that the cfDNA analysis integrates multiple metastatic tumor lesions. The whole exome-based analysis of this cancer revealed a profound spatial-temporal evolution of the tumor and the presence of multiple oncogenic and tumor suppressor gene alterations that could contribute to tumor progression, incomplete EGFR inhibitor response (i.e. disease persistence), and resistance to EGFR inhibitor treatment, even beyond the *EGFR^T790M^* mutation alone. The data indicated that the truncal mutations accounted for over 75% of the coding mutational burden at diagnosis but only 50-58% of the coding mutational burden at the time of full cancer evolution (patient death) in this case, with the emergence of subclonal mutations through tumor progression and both first- and second-line EGFR inhibitor resistance (Fig 1a). Overall, the findings in this case suggest an unanticipated parallel evolution of both the *EGFR^T790M^* mutation and two different *PIK3CA* driver alterations and are consistent with a genetic epistasis model in which the co-occurrence and co-evolution of several oncogene and tumor suppressor co-alterations present in the cancer influenced tumor progression and clinical outcome.

**Figure 1.**
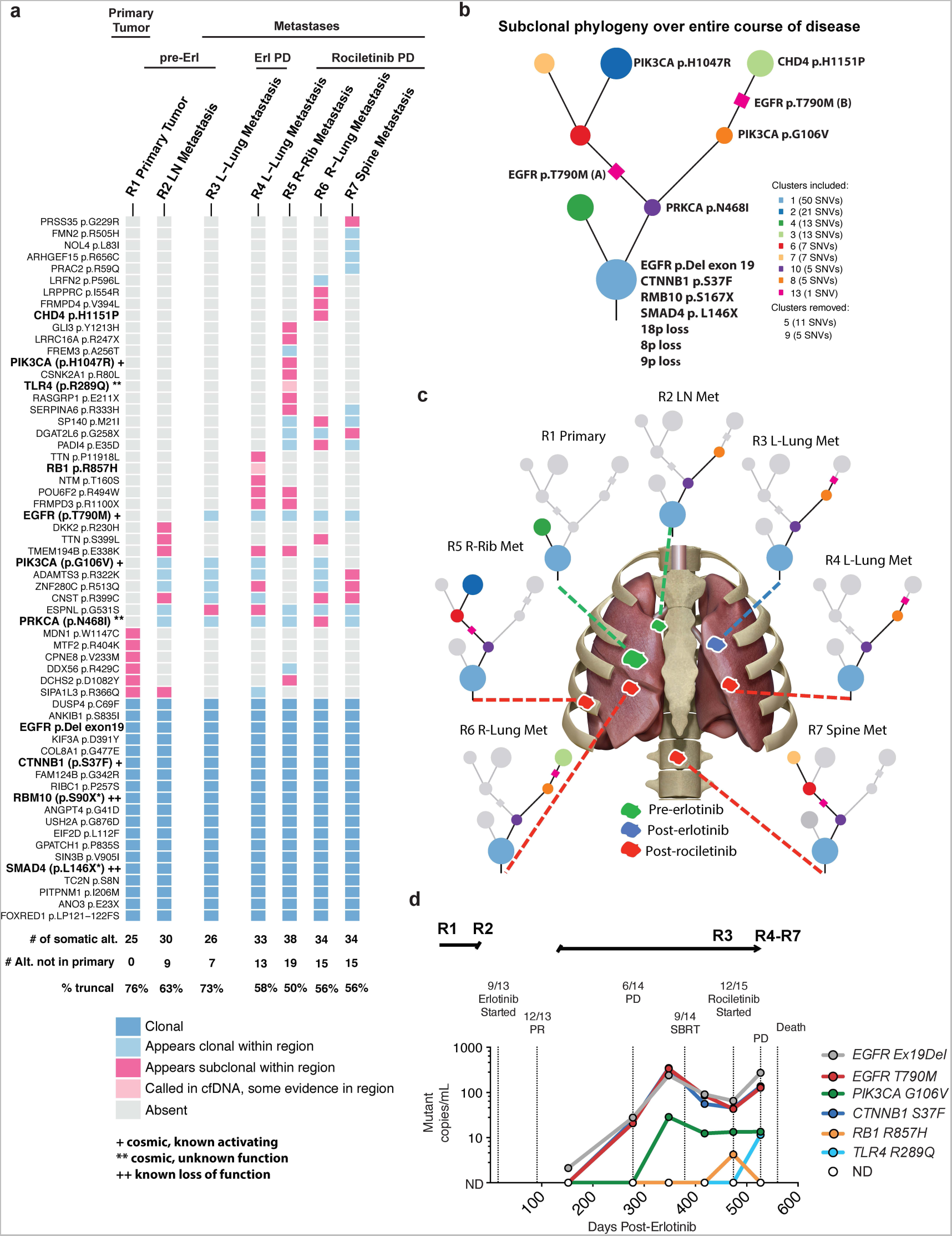
Comprehensive longitudinal genomic analysis of tumor and cell-free DNA of a patient with EGFR-mutant lung cancer from diagnosis to death. (a)Heatmap depicting the clonal status of non-synonymous somatic mutations including SNVs, dinucleotides and indels from each sequenced region of the patient’s disease as determined by subclonal copy number corrected cancer cell fraction and PyClone cross sample clustering. Somatic alterations were detected by whole-exome sequencing of the tumor DNA of the patient at initial presentation and surgical resection of EGFR-mutant lung cancer (R1), at the time of development of metastatic disease (R2), upon progression to first line treatment with erlotinib (R3), and at autopsy after treatment with the 2^nd^ line EGFR TKI rociletinib followed by PD and death (R4-R7). (see **Methods** for description of analysis). **(b)** Phylogenetic tree illustrating the evolutionary history of the patient’s disease at the level of subclonal clusters of mutations. These subclonal clusters are inferred, using PyClone, from the samples taken from the primary and different metastases at multiple time-points. The mutations were clustered based on their prevalence (subclonal copy number corrected cancer cell fraction) in the sequenced cancer cell populations across all samples, this clustering is then used to infer the founding clone (at the bottom of the tree) and subclonal clusters.. **(c)** Pictorial representation of primary tumor and metastatic sites analysed by whole exome sequencing. **(d)** ctDNA detectable in plasma from patient at indicated time points as determined by CAPP-Seq analysis.

The oncogenic alleles of both *CTNNB1* (β-catenin) and *PIK3CA* (G106V) were of particular functional interest in the genetic evolution of this cancer and we set out to define their potential roles, individually and in concert in the context of *EGFR-*mutant NSCLC. Oncogenic activation of *CTNNB1* has not been linked functionally to tumor progression or incomplete EGFR inhibitor response in EGFR-mutant NSCLC patients, while *PIK3CA^G106V^* has not previously been shown to impact NSCLC progression or EGFR inhibitor response. *PIK3CA^H1047R^* has been linked to lung cancer progression^22^ and promotes an invasive and metastatic phenotype when expressed in mammary epithelial cells^25^. The fact that *PIK3CA^H1047R^* was found at a metastatic site (R5) that did not harbor the *PIK3CA^G106V^* mutation (Fig. 1a-c) suggests that tumor metastasis is at least partially dependent on PI3K pathway activation. Functional analysis of CTNNB1 nuclear expression and the levels of phosphorylated AKT (as a measure of β-catenin and PI3K signaling activation, respectively) in the patient’s tumor samples confirmed activation of each pathway particularly in specimens with these activating alterations in *CTNNB1* and *PIK3CA* (Fig. S4). Additional functional studies revealed that expression of *CTNNB1^S37F^* in EGFR-mutant NSCLC cells limited EGFR inhibitor response by suppressing apoptosis, without significantly enhancing growth (Fig. S5a-c). These findings suggest that *CTNNB1^S37F^* may have promoted the incomplete response to EGFR inhibitor treatment (erlotinib, rociletinib) in the patient, offering evidence of a potentially clinically-relevant functional role for genetic epistasis in modulating treatment response. Expression of *CTNNB1^S37F^* also promoted cellular invasion, but not migration (Fig. S5d-e). In contrast, expression of *PIK3CA^G106V^* failed to limit response to EGFR inhibitor treatment, but instead promoted both invasion and migration (Fig. S5a-e). These data suggest that distinct evolutionary pressures selected for the presence and outgrowth of the *CTNNB1^S37F^* and *PIK3CA^G106V^* co-alterations, which may impart specific and non-redundant functions to promote either tumor progression (*PIK3CA^G106V^, CTNNB1^S37F^*) or limit EGFR inhibitor response (*CTNNB1^S37F^*). We found a trend towards the increased frequency of *CTNNB1^S37F^* in advanced-stage versus early-stage EGFR-mutant lung adenocarcinomas in the TCGA (The Cancer Genome Atlas) dataset (that is limited in the number of advanced-stage EGFR-mutant tumors) (Fig. S6), consistent with a role for *CTNNB1^S37F^* in metastatic EGFR-mutant NSCLC progression. Coupled with our functional data, these findings suggest β-catenin activation has multifactorial functions that promote tumor metastasis, consistent with prior preclinical studies^26^, and also contributes to incomplete EGFR inhibitor response. *PIK3CA^G106V^* was not identified in the primary tumor, but only emerged upon lymph node metastases and then persisted during both EGFR inhibitor treatments (Fig. 1a-b, S3). Our functional data suggest that PI3K pathway activation via *PIK3CA^G106V^* promoted metastatic progression, like *CTNNB1^S37F^,* but did not impact EGFR inhibitor response (unlike *CTNNB1^S37F^*), consistent with other recent clinical data^27^. These effects of *PIK3CA^G106V^* provide a unique example of the potential functional relevance of a subclonal driver in a treatment-naïve setting, extending prior reports that have instead generally implicated subclonal drivers in treatment resistance. Together, the findings provide evidence suggesting the biological and clinical relevance of cooperative polygenic alterations, i.e. genetic epistasis, in driving EGFR-mutant NSCLC pathogenesis and clinical progression. While the function of the *CHD4^H1151P^*mutation we detected is unknown, the function of CHD4 in chromatin remodeling^28^ suggests that it may play a role in epigenetic regulation of gene expression in this cancer, which has been linked previously with EGFR inhibitor tolerance^29^. The *TLR4^R289Q^* mutation we detected is predicted (*Mutation Assessor;cbioportal.org*) to impact function and may be an activating variant^30^ linked to NF-κB activity, which we have shown can promote EGFR inhibitor resistance^31,32^. Future studies beyond the scope here will be required to assess the functions of these additional genetic alterations.

These data highlighted the possibility that tumor progression and EGFR inhibitor response may be strongly influenced not only by genetic alterations in *EGFR* itself, but also by the genetic complement of concurrent alterations that are present and functional in each patient’s cancer. While the capture and polygenic analysis of tumor biopsies from large numbers of advanced-stage NSCLC patients with associated treatment response data remains challenging due to the difficulties in collecting multiple biopsies longitudinally from individual patients for tumor exome analysis, the advent of cfDNA profiling now makes it possible to more robustly study the link between tumor genetic alterations and treatment outcomes in this advanced-stage NSCLC disease context. To test whether genetic epistasis could impact response and resistance to EGFR targeted therapy more broadly in patients, we examined the polygenic landscape of gene alterations present in a substantial cohort of patients for whom longitudinal cfDNA analysis as well as clinical context and treatment response data were available (n=113 samples from 81 patients). The somatic genetic alterations present in these patients were identified using a clinically-validated plasma-based cfDNA exome analysis that measures single-nucleotide variants, small insertions/deletions, gene rearrangements/fusions, and amplifications across up to 70 clinically-relevant cancer genes (Table S4, S5, Methods)^33,34^. The use of this cfDNA assay, while not whole exome, offers the advantage of specifically determining whether co-alteration of many of the most clinically-important cancer-associated genes present in NSCLC (and other cancers) together with validated mutations in *EGFR* influences advanced-stage EGFR-mutant NSCLC patient clinical outcomes.

We analyzed 18 samples from patients obtained prior to initiating EGFR inhibitor treatment (erlotinib or afatinib), 11 samples from patients during either a partial response (PR) or stable disease (SD) to first-line EGFR inhibitor treatment (erlotinib), 46 samples from patients at the time of progressive disease (PD) to this first-line treatment (43 erlotinib, 2 afatinib, 1 erlotinib + bevacizumab), 16 samples from patients during a PR or SD to second-line treatment, and 22 samples from patients with PD on second-line treatment (chemotherapy n= 11, or EGFR inhibitor therapy n=11; 6-rociletinib, 2-osimertinib, 3-erlotinib/afatinib-based); 25 patients had more than one time-point analyzed (Fig. S7, Table S6). Somatic mutations were filtered to remove synonymous mutations as well as mutations of unknown significance, allowing focus on the most functionally relevant alterations (Methods). The number of detectable somatic alterations increased with each line of therapy, irrespective of age, gender, or tobacco exposure (pre-TKI: 2.2, PD 1^st^ line: 2.8, PD 2^nd^ line: 4.6, R^2^ = 0.1843, Slope 1.24, p < 0.0001; (Fig. 2b, S8). Enrichment for the *EGFR^T790M^* mutation was found upon progression on first-line EGFR inhibitor treatment (56.5% vs. 0%, p = 1.0E-5), as expected based upon the rare detection (~0.5%) of *EGFR^T790M^* prior to first-generation EGFR inhibitor treatment^35^ and the established incidence of *EGFR^T790M^* (55%-65%) upon progression on first-generation EGFR inhibitors^21^ (Fig. 2a,b, Table S6). Upon progression on second-line treatment (EGFR inhibitor or chemotherapy), the analysis revealed further selection for co-alterations in *CCNE1* (18% vs. 2.1%, p = 0.03), and *KIT* (13.6% vs. 0%, p = 0.03), with trends towards increases in alterations in *MYC* (18% vs. 4%, p = 0.08), *PIK3CA* (23% vs. 8.6 %, p = 0.14), and *CTNNB1* (18% vs. 10%, p = 0.11) (Fig. 2c). Alterations in genes involved in receptor tyrosine kinase (RTK), MAPK, Cell Cycle, PI3K, and WNT signaling were more frequently found in patients who had progressed on 2^nd^ line therapy (Fig. 2d and Table S6). The mean number of functional alterations detected in cfDNA was lower in patients who responded to subsequent EGFR inhibitor therapy versus those who did not respond to this subsequent treatment (2.3 vs. 3.9, p=0.0005, Fig. S9, Table S7), with patients harboring gene level *MET* (p=0.03) or *TP53* (p=0.06) alterations being least likely to respond to subsequent EGFR inhibitor treatment (Fig. S9). Pathway level TP53 alterations also correlated with non-response to EGFR inhibitor treatment (27% responders vs. 58% non-responders, p=0.03) as did alterations in cell cycle genes (0% responders vs. 25% non-responders, p=0.004) (Fig. S9). In the context of the cfDNA targeted exome profiled, these data suggest further selection for increased genetic diversity during iterative tumor progression on therapy. The data provide evidence in support of a model in which the polygenic landscape of *EGFR*-mutant NSCLC may contribute to clinical progression on EGFR inhibitor treatment, in a manner that augments or complements the effects of *EGFR* gene alterations (e.g. *EGFR^T790M^*).

**Figure 2.**
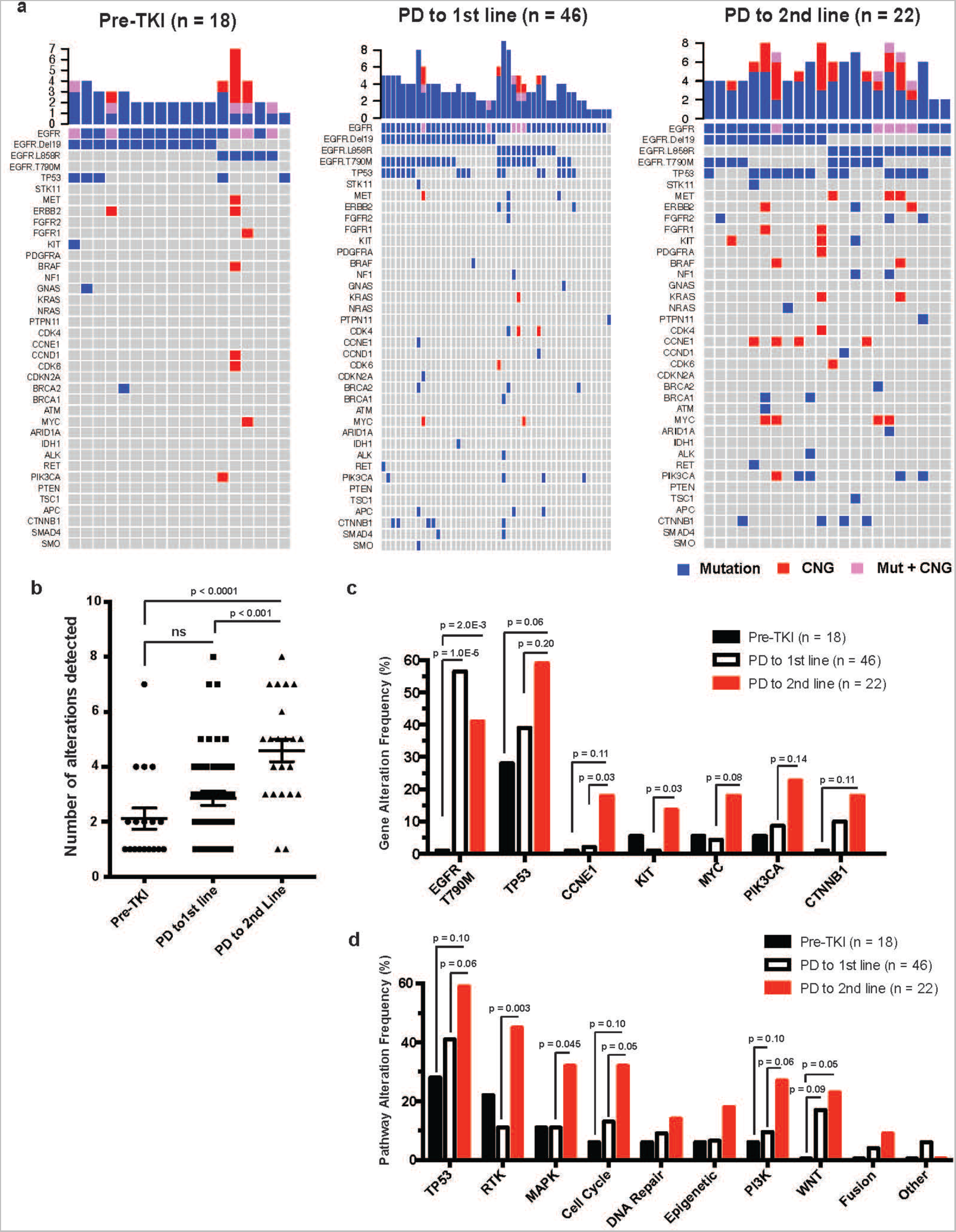
Cancer therapy-induced evolution of genomic alterations detected in cell-free DNA of advanced-stage EGFR-mutant NSCLC patients. (a) Circulating tumor DNA analysis of 86 samples collected from 81 patients with known clinical history. Samples were segregated based on whether they were collected prior to EGFR TKI treatment; pre-TKI (n=18), at the time of progression to first-line EGFR TKI therapy; PD to 1^st^ line (n=46), or at the time of progression to 2^nd^ line anti-cancer therapy (2^nd^ or 3^rd^ generation EGFR TKI, or chemotherapy); PD to 2^nd^ line (n = 22). Non-synonymous mutations with predicted functional impact and copy number gains (CNG) are indicated. **(b)** Number of functional alterations detectable based on line of therapy are indicated. P-value determined by ANOVA test with Tukey correction for multiple comparisons. **(c)** Changes in gene alteration frequency with line of therapy **(d)** Changes in cancer-related pathway alterations (as defined in **Table S5**) with line of therapy. **(b and c)** Two-way Fisher’s exact test was performed to identify statistically significant differences between pre-TKI and PD to 1^st^ line, between PD to 1^st^ line and PD to 2^nd^ line, and between pre-TKI and PD to 2^nd^ line.

Consistent with this notion, the analysis of several individual cases of EGFR-mutant NSCLC for which longitudinal cfDNA profiling was available (Fig. 3a-e, S10) showed that progression on the third-generation EGFR^T790M^ inhibitors osimertinib and rociletinib occurred during co-selection for the simultaneous emergence of multiple oncogenic mutations. *CTNNB1*^S37F^ plus *CDK4, FGFR1, KIT, KRAS,* and *PDGFRA* CNGs emerged in one case of acquired rociletinib resistance (Fig. 3a). Primary osimertinib resistance was evident in a patient with co-occurring mutations in *TP53^V272M^, PIK3CA^D549Y^,* and *CCNE1* CNG (Fig. 3b). In another case, the emergence of erlotinib resistance and then primary resistance to osimertinib was associated with co-occurrence of *EGFR^T790M^, CCND1^E9D^, CDH1^D400N^* mutations and CNGs in *CDK4, EGFR,* and *KRAS* (Fig. 3c). Acquired osimertinib resistance emerged in the setting of co-occurring *EGFR^C797S^, CTNNB1*^S37F^, and a *SMAD4^Q516^*^*^ loss-of-function mutation (Fig 3d). Co-emergence of *TP53^C277F^* along with CNGs in *BRAF, CCNE1, EGFR, MYC,* and *PIK3CA* occurred during rociletinib resistance in another patient (Fig 3e), who subsequently developed *MET^H1112Y^,* along with *MET* and *FGFR1* CNGs upon resistance to osimertinib (Fig. 3e). Concurrent alterations including a *PIK3CA^E545K^* activating mutation along with loss-of-function co-mutations in *TP53^exon11Del^* and *PTEN^S287*^* were seen in another patient with acquired resistance to osimertinib (Fig. S10). Treatment with multiple EGFR-directed targeted therapy regimens drove selection for multiple concurrent alterations in BRAF(V600E), ERBB2(R143Q) and MET(CNG) at distinct time points during the course of treatment and iterative tumor progression in another case (Fig. S10).

**Figure 3.**
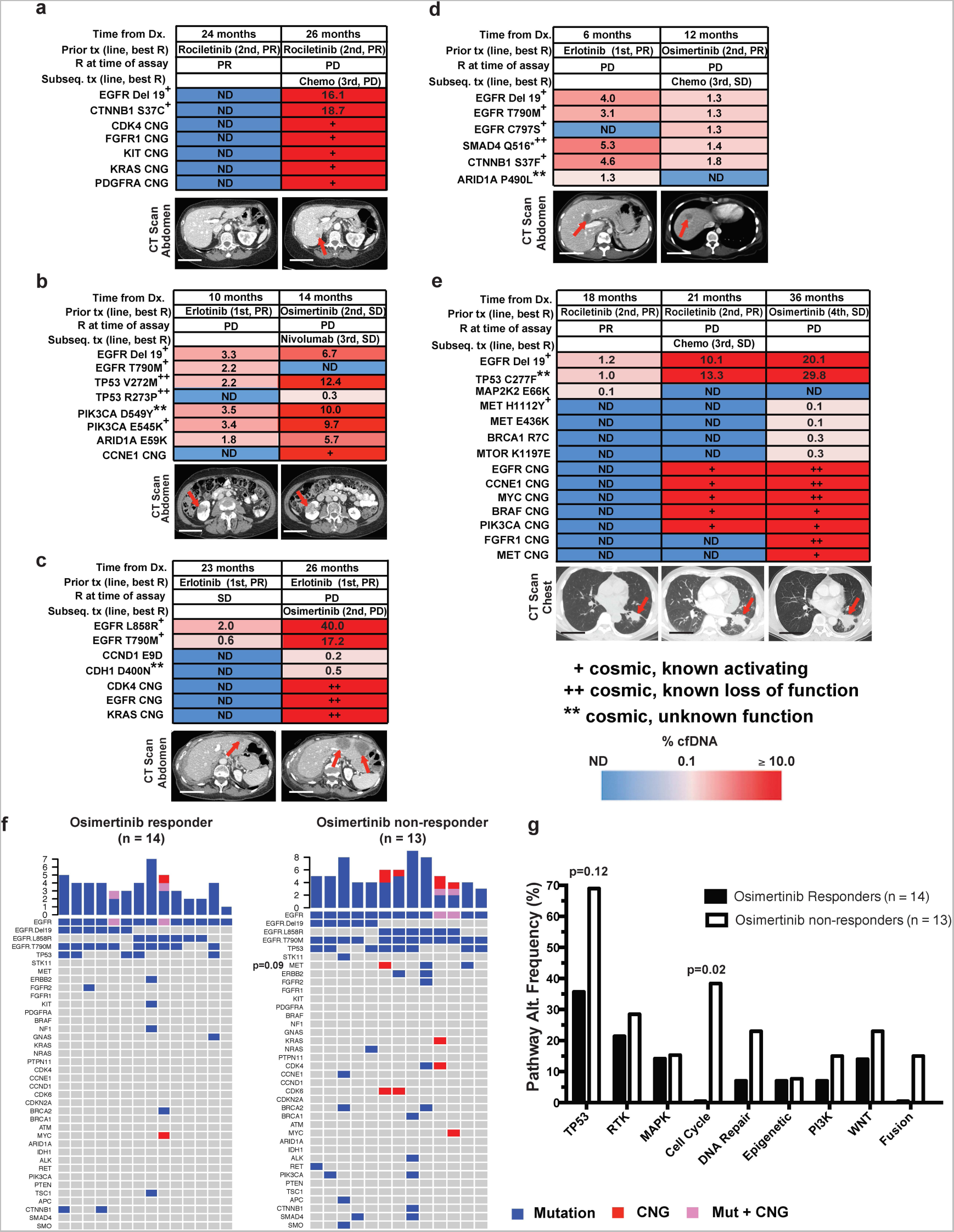
Patient level evolution of genomic alterations detected in cell-free DNA of advanced-stage EGFR-mutant NSCLC patients. (a-e) Circulating tumor DNA (ctDNA) alterations detectable by clinical assay in patients with EGFR mutations. Mutant allele frequency of detected ctDNA alterations are indicated, as is line of therapy and clinical response pre- and post-assay. Computed-tomography (CT) scans of the chest and abdomen are shown, demonstrating sites of metastases (red arrows). Scale bar = 5 cm, ND (no alteration detected). PR (Partial Response), PD (Progressive Disease), SD (Stable Disease). Plasma copy number gain (CNG) of 2.0-2.49 is reported as **+** and 2.5-4.0 as **++ (Methods)**. **(f)** Genomic alterations detectable in ctDNA of patients who were subsequently treated with osimertinib and exhibited a radiographic response (left, demonstrated PR on imaging by clinician assessment) versus patients who did not respond (right, SD or PD on radiographic imaging by clinician assessment, or clinical progression leading to death prior to imaging being performed) versus patients who did not respond (right, by clinician assessment as defined above). **(g)** Pathway alterations (defined in **Table S5**) in osimertinib responders vs. non-responders demonstrated increased cell cycle gene alterations in non-responders. (f-g) P-values determined by two-tailed Fisher’s Exact test.

Our data suggested a potential role for genetic epistasis in controlling tumor progression and EGFR inhibitor response in EGFR-mutant NSCLC patients. If this hypothesis is true, then the presence of specific cooccurring genetic alterations would be enriched in non-responder versus responder patients. Primary resistance of *EGFR^T790M^*-positive NSCLCs to initial osimertinib treatment was associated with the presence of alterations in cell cycle genes such as *CDK4/6* (5/13, 38% vs. 0/14, 0%, p = 0.02), and a trend towards alterations in *MET* (3/13, 23% vs. 0/14, 0%, p = 0.09) and *TP53* (8/13, 69% vs. 5/14, 36%, p=0.12) prior to treatment (Fig. 3f-g, Table S8). While mechanisms of acquired osimertinib resistance have been reported^36-38^, these data uncover potential clinically-relevant mechanisms that may limit the initial response to osimertinib treatment (i.e. promote primary resistance). The findings suggest that multiple co-alterations in specific signaling pathways that promote biological processes critical for cancer growth may help limit EGFR inhibitor response, even in cancers with *EGFR^T790M^.* This finding of polygenic driver alterations in the *EGFR^T790M^-*positive cases may help explain why clinical responses to EGFR^T790M^-directed therapies such as osimertinib occur in only 60-70% of *EGFR^T790M^*-positive NSCLC patients and are almost always incomplete^4,5^. Altogether, the data argue that EGFR-mutant NSCLC clinical outcomes are impacted by the widespread presence of combinations of functional genetic alterations that are present and undergo selection before and during EGFR inhibitor treatment and tumor progression.

To provide further support for this genetic epistasis model, we undertook a large-scale analysis using the clinically-validated cfDNA assay (Table S4, S5, Methods) to test the hypothesis that concurrent oncogenic alterations might be a prevalent feature of advanced-stage EGFR-mutant NSCLC more broadly. The clinical cohort we profiled consisted of 1150 *EGFR* mutation positive and 1008 *EGFR* mutation negative patients with advanced-stage (Stage III and IV) NSCLC (Table S9), of which a subset was analyzed above (Fig. 2 and 3). To enhance focus on the most functionally relevant genetic alterations, we filtered for mutations that were non-synonymous and validated or predicted to impact gene function (Methods), which yielded 1150 EGFR-mutation-positive and 945 EGFR mutation-negative cases. While clinical outcomes data are not readily available for most patients represented in this larger sample set, this cfDNA cohort contains advanced-stage NSCLC patients who are often heavily pre-treated and as such it differs from TCGA and other genomic compendia of lung cancer that are based largely on the analysis of early-stage resected tumors in patients who generally are not as exposed to systemic drugs such as EGFR inhibitors. Therefore, this cfDNA cohort represents the landscape of resistance and the initial driver/founder mutations. Analysis of the 1150 EGFR-mutant patient cohort revealed the widespread presence of concurrent genetic alterations, in addition to the *EGFR* driver mutation (Fig. 4a). Overall, the vast majority of patients (93.1%; 1071 out of 1150) harbored at least one additional variant of known or likely functional significance (Table S10). The EGFR-mutant cases contained an average of 3.6 genetic alterations (out of the ≤70 genes profiled), including the *EGFR* mutation that was present (range 1-18). There was enrichment for co-alterations in *CTNNB1* (5.3% vs. 1.8%, p = 2.2E^−05^), *CDK6* (6.9% vs. 3.2%, p = 1.5E^−04^), *AR* (5.0% vs. 2.6%, p = 0.007), and a modest difference in *TP53* (55% vs. 50%, p = 0.04) in the EGFR-mutant cohort, compared to the stage-matched EGFR mutation-negative cohort similarly profiled using the cfDNA assay (n=945) (Fig. 4a-b, S11, S12, Table S10, S11). The pathway-level analysis showed selection for genetic co-alterations in *WNT/CTNNB1* and hormone signaling genes in the EGFR-mutant cohort, whereas alterations in several functional groups including RTKs and MAPK pathway genes (e.g. *KRAS*) were more enriched in the EGFR mutation-negative cohort (as expected) (Fig. 4c and S11). This large-scale dataset further suggests an important role for WNT/CTNNB1 signaling, consistent with a small case series^24^, and cell cycle gene aberrations in the pathogenesis of EGFR-mutant NSCLC, re-enforcing our earlier findings linking alterations in these genes and pathways to clinical outcomes and treatment response in EGFR-mutant NSCLC patients (Fig. 1-3). The majority (89.4%) of the genetic co-mutations present in the *EGFR* mutation-positive cohort have verified or likely functional impact (by *in silico* modeling, Methods, Table S10), with only 10.6% (370/3507) of these co-mutations classified as likely passenger events (neutral or unknown functional impact). In contrast, 16.4% (431/2622) of the mutations present in the *EGFR* mutation-negative cohort were classified as passenger events (p<0.0001 by Fisher’s exact test) (Table S11).

**Figure 4.**
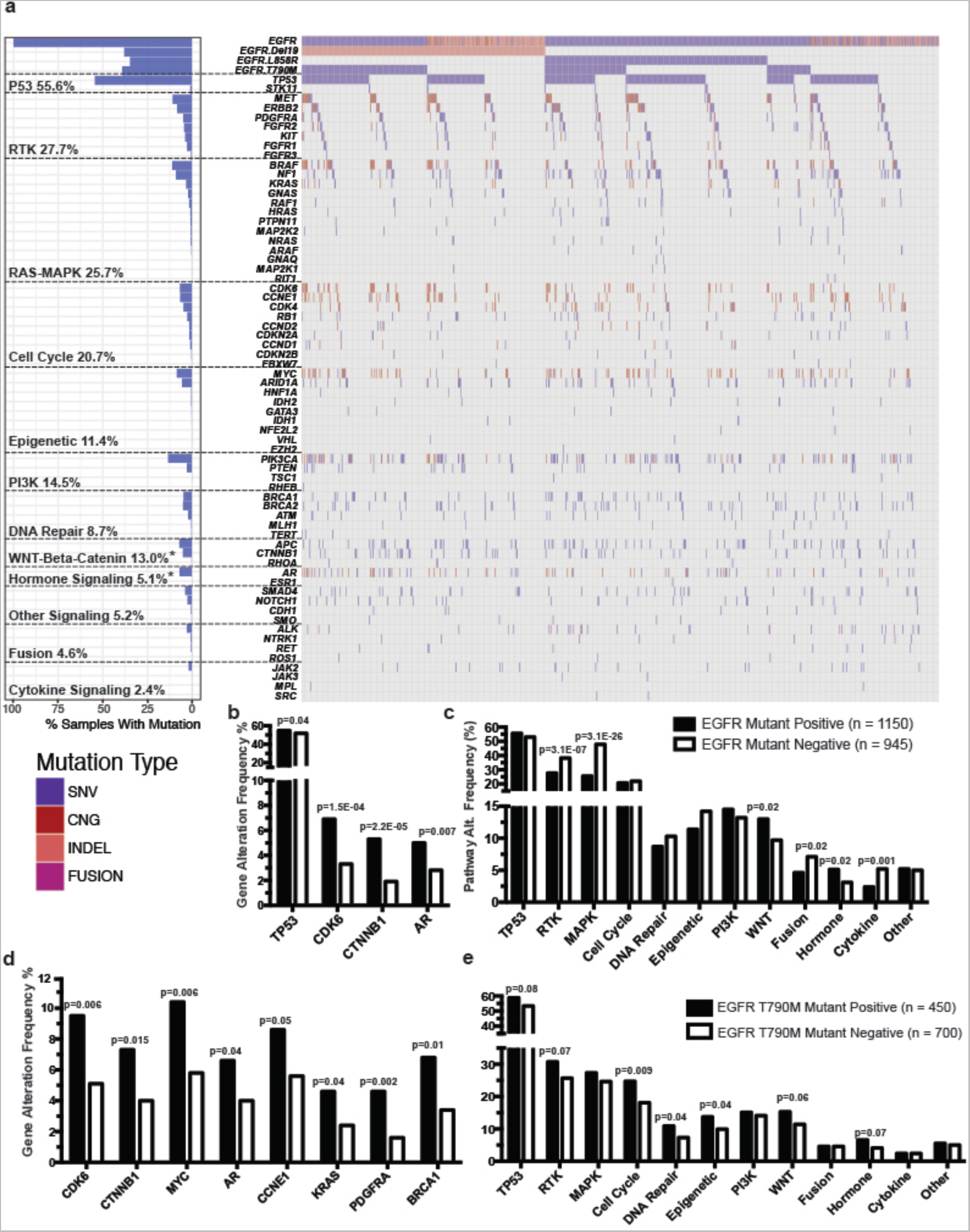
Concurrent genomic alterations detectable in cell-free DNA of EGFR-mutant positive compared to EGFR mutant negative non-small cell lung cancer (NSCLC) patients. **(a)** Frequency of genomic alterations: non-synonymous somatic variants of predicted functional significance (SNV, see Methods), copy number gains (CNG), insertions or deletions (INDEL), or gene rearrangements (FUSION) in the cancer-related genes listed, detected by next-generation sequencing of circulating tumor DNA from 1150 advanced-stage EGFR-mutant positive NSCLC patients. The frequency of alterations in a given gene were compared to a cohort of 945 EGFR-mutant negative NSCLC patients in **Supplementary Figure 11. (b)** Gene alterations detectable in the cell-free DNA of 1150 advanced EGFR-mutant positive compared to 945 EGFR-mutant negative patients (Two-tailed Fisher's exact test performed to identify statistically significant differences in *TP53, CDK6, CTNNB1,* and *AR.* **(c)** Differences in pathway level alterations between EGFR-mutant positive and EGFr-mutant negative cases by Fisher's Exact test comparing EGFR-mutant positive to EGFR mutant-negative. **(d-e)** Gene and pathway level alterations detectable in the cell-free DNA of 700 advanced EGFR-mutant, EGFR T790M negative compared to 450 EGFR-mutant T790M positive patients (Two-tailed Fisher's exact test performed to identify statistically significant differences) based on the findings in **Supplementary Figure 13**.

Our earlier data (Figs. 1-3) suggest that genetic collectives and epistasis, rather than only mutation of *EGFR,* influences tumor progression and clinical outcomes. This model predicts that alterations in specific genes or pathways beyond EGFR would occur during the emergence of EGFR inhibitor resistance in patients (e.g. beyond the presence of a resistance-conferring mutation in *EGFR* itself). As in Figs. 1-3, the analysis of EGFR inhibitor-treated patients can provide evidence that concurrent alterations are functional, as those that are not selected for (i.e. passenger alterations) are unlikely to undergo recurrent selection across a large cohort of patients such as the one presented here. The *EGFR^T790M^* mutation arises in over 50% of cases of acquired resistance to first-generation EGFR inhibitors (erlotinib, gefitinib)^21^ while very rarely being detectably present prior to EGFR inhibitor treatment^35^, providing an opportunity to test this hypothesis. In the cohort of 1150 EGFR-mutant NSCLC samples that we analyzed, 450 had a detectable *EGFR^T790M^* mutation. While EGFR inhibitor response data are not available for most of the patients in this large cohort, it is likely that the vast majority of these *EGFR^T790M^*-positive patients had been treated with an EGFR inhibitor based on the scarcity of *EGFR^T790M^* mutations detectable prior to treatment (0.5%)^35^. Analysis of the genetic co-alterations present in the 450 *EGFR^T790M^*-positive cancers revealed more frequent alterations in certain pathways compared with the *EGFR^T790M^*-negative cases (n=700) (Fig. 4d-e, S13, Table S12 and S13). This specific enrichment included co-alterations in genes and pathways controlling the cell cycle (*CDK6* and *CCNE1* CNGs), WNT signaling (*CTNNB1* oncogenic mutations), hormone signaling (androgen receptor, *AR,* somatic mutations), epigenetic modification *(MYC* CNG), *KRAS* and *PDGFRA* gene alterations (CNG and oncogenic mutations), as well as *BRCA1* mutations being more common in the *EGFR^T790M^*-positive cases (Fig. 4d, S14, S15). In a sub-analysis of *EGFR^C797S^* mutation-positive cases (n=15), which are predicted to arise upon acquired resistance to osimertinib^38^, recurrent activating alterations in MAPK pathway (including *KRAS* CNG and oncogenic mutations) and cell cycle genes (*CDK4, CDK6*), and *AR* CNGs were identified (Fig. S16). These data further suggest that multiple co-alterations in specific genes and pathways controlling cancer growth, likely in cooperative or non-redundant ways may contribute to incomplete EGFR inhibitor response and resistance, even in cancers with *EGFR^T790M^* and *EGFR^C797S^.* This finding of polygenic oncogenic alterations in the larger cohort of *EGFR^T790M^*-positive patients is consistent with our earlier clinical data linking several of these genes and pathways to clinical outcomes and treatment response in EGFR-mutant NSCLC patients (Fig. 1-3). Together, these data in EGFR^*T790M*^-positive cases offer insight into the potential basis of the typically incomplete clinical responses to EGFR^T790M^-directed therapies such as osimertinib in *EGFR^T790M^*-positive NSCLC patients and identify potential combination therapy strategies to enhance response.

By extension of our initial tumor biopsy exome analysis (Fig. 1), we assessed whether these genetic co-alterations detected in the cfDNA were clonal versus subclonal in EGFR-mutant NSCLC. A method for assessing clonality in tumor cfDNA has not been established, in contrast to the analysis of tumor samples by exome deep sequencing. Our strategy was to compare the mutant allele frequency (MAF) for an altered gene to the maximum MAF detected in a given sample. To estimate the relative proportions of cfDNA that would differentiate a mutant allele as clonal versus subclonal, as a benchmark we correlated the tumor-based clonality analysis in Fig. 1a with the cfDNA mutational analysis from the same patient upon progression on rociletinib (Fig. 1c). The *CTNNB1^S37F^* mutation was clonal in the patient tumors at autopsy (Fig. 1a and S3, S4), but present at only 50% of the maximum mutant allele (*EGFR^ex19del^*) detected in the cfDNA (Fig. 1c). In contrast, the *PIK3CA^G106V^* mutation was subclonal (only present in 2 of 4 metastatic sites) at autopsy (Fig. 1a,b and S3) and correspondingly present at only 5% of the maximum MAF in the cfDNA (Fig. 1d). Based on this analysis, we estimated that clonal alterations may be detectable as low as 20% of the maximum MAF in ctDNA, but likely to be subclonal when detected at < 20%. Using this approach, we found that *EGFR* mutation-positive NSCLCs are more likely to harbor subclonal genetic alterations than *EGFR* mutation-negative NSCLCs (Tables S10, S11,Fig. S17a, 34.1% subclonal alterations in EGFR-mutation positive vs. 26.6% subclonal events in EGFR-mutation negative cases, p = 1.759e^−9^). Subclonal alterations were also more commonly found in tumors that harbored *EGFR^T790M^* (Tables S12, S13, Fig. S17b, 37.2% in *EGFR^T790M^-* positive vs. 31.3% in *EGFR^T790M^*-negative cases, p = 4.531e^−04^). As expected, founder canonical *EGFR* mutations (L858R, exon19del) were most often estimated as clonal (~96.3%, 386/401; ~95%, 419/439; respectively). Relative to the clonal founder *EGFR* mutation, the *EGFR^T790M^* resistance mutation was more frequently subclonal (~75% clonal; 338/450, p < 0.0001 compared to canonical EGFR mutations, L858R or exon19del), a finding of clinical relevance given that subclonal *EGFR^T790M^* may be linked to inferior clinical response to third-generation EGFR inhibitor treatment^39^. *EGFR^C797S^,* which causes acquired osimertinib resistance^38^, was also frequently subclonal (47% (7/15) clonal; p = 0.0295 compared to *EGFR^T790M^*). Co-alterations in *CTNNB1* (75%, 48/64), *TP53* (72.9%, 562/771), and *PIK3CA* (71.8%, 79/110) were frequently clonal, while *AR* mutations were more frequently subclonal (clonal in 38%, 8/21) of EGFR-mutant samples. These observations are consistent with our earlier findings (Fig. 1-3) in support of a genetic epistasis model wherein both clonal and subclonal evolution of several oncogene and tumor suppressor co-alterations, beyond *EGFR* mutations alone, may influence EGFR-mutant NSCLC progression and drug response. Overall, the data uncover a previously underappreciated degree of selection for specific genetic co-alterations in cancer-relevant genes within advanced-stage EGFR-mutant NSCLCs, providing evidence that the pathogenesis of this subtype of NSCLC may be polygenic and not merely monogenic (i.e. controlled by mutant EGFR alone).

The uncommon opportunity to longitudinally capture multiple tumor biopsies from an individual patient over several years: from the time of the initial diagnosis of early-stage disease, to metastatic progression, to both first- and second-line treatment progression, and finally death, and analyze them on a whole exome-wide level in coordination with cfDNA profiling of this case as well as a substantial series of additional patients has provided a window into the diverse and evolving genetic architecture of *EGFR*-mutant NSCLC. To our knowledge, the clinical case we present is one of the more extensive analyses of coordinated longitudinal tumor and cfDNA exome profiling in NSCLC that has been linked to clinical outcome data obtained over many years of disease progression. Our data offer support for a model that stands in contrast to the prevailing view (Fig. S18). Instead of classifying genetic subtypes of advanced-stage NSCLC discretely as monogenic diseases (e.g. EGFR-mutant NSCLC) and treating with monotherapies (e.g. an EGFR inhibitor), our data argue for an important function of genetic epistasis in promoting specific, non-redundant tumor-driving phenotypes, such as invasion/metastasis or incomplete therapy response and resistance, reminiscent with recent findings in myeloproliferative neoplasms^40^. Our diverse datasets highlight new context-specific cooperative roles in cancer pathogenesis for co-alterations in important cancer genes such as β-catenin, *PIK3CA, MYC* and several cell cycle genes (e.g. *CDK4/6*) in driving EGFR-mutant NSCLC progression. The data also highlight the notion that critical pathways that drive either tumor progression or resistance to therapy can evolve at multiple sites in parallel within the same patient, and likely within distinct tumor cell subpopulations. Moreover, our unanticipated finding that cell cycle gene aberrations (e.g. *CDK4/6* and *CCNE1* amplifications) are absent in responders to EGFR inhibitor therapy, but present in 30-40% of non-responders (including to the third-generation EGFR inhibitor osimertinib) warrants further investigation. Interestingly, preclinical NSCLC models have shown synergy between EGFR and CDK4/6 co-inhibition (with patient-ready drugs including the FDA-approved CDK4/6 inhibitor palbociclib)^41,42^. Together, these prior findings and our clinical and genetic data suggest that a clinical trial assessing the combination of EGFR and CDK4/6 inhibition in EGFR-mutant NSCLC patients with evidence of tumor genomic cell cycle alterations (e.g. *CDK4/6* CNG) should be considered. Our findings provide a roadmap for the future elucidation of the specific functional roles and interactions of the broader set of genetic co-alterations comprising the polygenic landscape of advanced-stage EGFR-mutant NSCLC that we uncover here, paving the way for additional potential polytherapy trials to enhance the magnitude and duration of response in patients.

Our findings prompt a re-consideration of advanced-stage EGFR-mutant NSCLC as largely a polygenic rather than monogenic disease, thus re-shaping the understanding of the basis of NSCLC established by earlier important genomic studies that focused predominantly on early-stage lung cancer (e.g. TCGA) in patients not receiving systemic targeted therapy. Our data highlight the importance of deploying more informed molecular diagnosis, monitoring, and dynamically-applied rational polytherapy strategies to address the genetic epistasis of clonal and subclonal alterations that our data reveal in order to better control this deadly cancer.

## Methods

### Whole-exome sequencing and analysis

Informed consent was obtained from the patient and patient’s family for study of biological materials and clinical records obtained from the patient. DNA was extracted from FFPE for primary tumor and frozen tumor tissue samples and matched non-tumor tissue using the Qiagen Allprep DNA/RNA Mini Kit. The library preparation protocol was based on the Agilent SureSelect Library Prep and Capture System (Agilent Technologies, Santa Clara, CA). Quantitation and quality were assessed using the Qubit Flourometer (Thermo Fisher). DNA concentration was determined to be greater than 2.5 ng/ul and the overall quantity > 500ng. By Nanodrop, the 260/280 ratio was greater than 1.7. DNA was resuspended in a low TE buffer and sheared (Duty Cycle 5%; Intensity 175; Cycles/Burst: 200; Time: 300s, Corvaris S2 Utrasonicator). Bar-coded exome libraries were prepared using the Agilent Sure Select V5 library kit per manfucaturer’s specifications. The libraries were run on the HiSeq2500.

### Alignment

Raw paired end reads (100bp) in FastQ format generated by the Illumina pipeline were aligned to the full hg19 genomic assembly obtained from USCS, gencode 14, using bwa version 0.7.12. Picard tools version 1.117 was used to sort, remove duplicate reads and generate QC statistics. Tumor DNA was sequenced to median depth of 303X (range 114.39-383.41) and the matched germline DNA to average depth of 231.65.

### Exome analysis

SNV, INDEL and Dinucleotide substitution calling, identification and classification of driver mutations, somatic copy number aberration calling, subclonal deconstruction and phylogenetic tree construction were performed as described in [*Jamal-Hanjani et al. TRACERx – Tracking Non-Small Cell Lung Cancer Evolution, 2017 (in press)]*].

### Classification of SCNAs

SCNA events were defined as segments called by ASCAT >= 400kb in size that met set thresholds. Segments with a combined raw nMinor and nMajor greater than a 1.5 times the ASCAT derived ploidy for their specific tumor region were considered SCNA gains. SCNA losses had an integer nMinor value of 0 and a combined raw nMinor and nMajor of less than 1.25 times ploidy for their specific tumor region.

### Incorporation of T790M mutation into phylogenetic reconstruction

In order to create and accurate subclonal phylogeny it is necessary to remove mutation clusters that violate two evolutionary principles. Firstly the pigeonhole principle which ensures that two mutation clusters cannot be considered to be on separate branches of an evolutionary tree and thus be independent if the cancer cell fraction values of the two clusters together exceeds 100% within region of a tumor. Secondly, a descendent clone must have a smaller cancer cell fraction than its ancestor within each and every tumor region, referred to as the ‘crossing rule’. Using these principles it can be determined whether particular mutation clusters conflict with each other and cannot be fitted to the same evolutionary tree.

The subclonal phylogeny illustrating the entire course of the patient’s disease was derived following these two principles and the methods of multi-sample subclonal deconstruction and tree construction in [*Jamal-Hanjani et al. TRACERx – Tracking Non-Small Cell Lung Cancer Evolution, 2017 (in press)]*]. However, the EGFR T790M SNV did not cluster with any other SNVs following these methods due to its unique CCF profile across R3, R4, R5, R6 and R7. No other SNV appears clonal in all these regions as well as being absent from both R1 and R2. As cluster 7 and the EGFR T790M mutation appear clonal in R3, R4, and R6 but cluster 7 is absent from R5 and R7 and EGFR T790M is present they cannot, by the crossing rule, be present in the same population of cells. In addition, as cluster 7 was present clonally in R2 before Erlotinib treatment while T790M is absent from R2, it follows that cluster 7 is likely to have arisen before the T790M SNV.

The most parsimonious solution to this violation of the crossing rule, assuming that the cancer cell fractions are correct, is that there are two independent origins of T790M. T790M SNV (A) would occur in a cell already containing the SNVs from cluster 7, and go on to become clonal post-Erlotinib treatment in R3, R4 and R6. T790M SNV (B) would occur in a population of cells lacking the SNVs present in cluster 7 and go on to become clonal in R5 and R7 post-Erlotinib. These possible origins of the T790M are indicated on the subclonal phylogeny that can be seen in Figure X by the placement of a magenta square on the relevant branches.

### Data Availability

All exome sequencing data will be available through the European Genome-phenome Archive (EGA), accession number: Pending.

### Code availability

Most bioinformatics tools used in the analysis of this dataset are publicly available; any that are not are available on request.

### Cell-Free DNA Analysis

1150 consecutive EGFR-mutant positive advanced (stage III or IV) NSCLC samples from 1027 patients were collected and analyzed between June 2014 and April 2016. 1008 *EGFR*-mutant negative advanced (stage III or IV) NSCLC samples from 999 patients were collected and analyzed between January 2016 and April 2016. Samples were shipped to a Clinical Laboratory Improvement Act (CLIA)-certified, College of American Pathologists-accredited laboratory (Guardant Health, Redwood City, California). Cell-free DNA (cfDNA) was extracted from whole blood collected in 10mL Streck tubes. After double ultracentrifugation, 5ng - 30ng of cfDNA was isolated for digital sequencing as previously described ^33,43^. For EGFR-mutant positive NSCLC samples 28 were run on the 54-gene panel, 419 on the 68-gene panel, and 703 on the 70-gene panel (Table S4). All 1008 EGFR-mutant negative samples were run on the 70 gene-panel. Sequencing data was analyzed using a custom bioinformatics pipeline to identify single nucleotide variants (SNVs) in 70 genes (150kb panel footprint), CNGs in 18, indels in three (*EGFR* and *ERBB2* exons 19 and 20; *MET* exon 14), and *ALK, RET, ROS1, NTRK1, FGFR2,* and *FGFR3* fusions ^33,43^. All cell-free DNA fragments, both leukocyte- and tumor-derived, were simultaneously sequenced. The variant allele fraction (VAF) was calculated as the proportion of cfDNA harboring the variant in a background of wild-type cell-free DNA. The analytic sensitivity reaches detection of 1-2 mutant fragments in a 10 ml blood sample (0.1% limit of detection) with analytic specificity > 99.9999%^33,43^. CNGs were reported as the absolute gene copy number in plasma. For longitudinal case (Fig. 4d), cell-free DNA was isolated from 1 ml of frozen plasma and analyzed as described^17,44^. Clinical data was collected by review of medical records under an IRB-approved protocol (UCSF). Non-synonymous mutations from EGFR-mutant positive and negative datasets were further processed using R statistical computing program (version 3.3). Unknown significant variants were filtered out by using COSMIC (V79), GENIE(http://www.aacr.org/Research/Research/Pages/aacr-project-genie.aspx#.WMeBiRLytaM) and prediction algorithm (http://mutationassessor.org/r3/). For clonality analysis, the ratio of MAF over Maximum-percentage detection (Max-pct) were computed, the probability distribution was plotted using kernel density estimation. The value of 0.2 was defined as a robust cutoff for subclonal or clonal mutations derived from the case analysis in this paper. Two-tailed Fischer’s Exact test was used to determine statistically significant differences in ctDNA alterations between cohorts.

### Cell Lines and Reagents

The HCC827 (*EGFR* exon 19 deletion) and HEK293-FT cell lines were obtained, authenticated, and cultured as recommended by the American Type Culture Collection (ATCC). These cell lines confirmed to be negative for mycoplasma. HCC827 cells were cultured in RPMI 1640 media (Hyclone, GE Healthcare) supplemented with 10% FBS (SAFC, Sigma-Aldrich), 1X penicillin and streptomycin (UCSF, Cell Culture Facility). HEK293-FT cells were cultured in DMEM media (Hyclone, GE Healthcare), supplemented with 10% FBS, 0.1X penicillin and streptomycin. All cell lines were grown at 37 °C, in a humidified atmosphere with 5% CO2. Erlotinib and rociletinib were purchased from Selleckchem.

Mammalian expression vectors pQCXIB empty (w335-1) was a gift from Eric Campeau (Addgene plasmid # 17487)^45^; pBABE-puro was a gift from Hartmut Land & Jay Morgenstern & Bob Weinberg (Addgene plasmid # 1764)^46^; pCMV-VSV-G (Addgene plasmid # 8454) and pUMVC (Addgene plasmid # 8449)^47^ were a gift from Bob Weinberg; pBabe puro HA PIK3CA was a gift from Jean Zhao (Addgene plasmid # 12522)^48^; human beta-catenin pcDNA3 was a gift from Eric Fearon (Addgene plasmid # 16828)^49^. The PIK3CA and b-catenin constructs were engineered to express G106V *PIK3CA* and S37F *CTNNB1* following QuickChange II XL Site-Directed Mutagenesis Kit protocol (Agilent Technologies). The S37F *CTNNB1* fragment was then sub-cloned in a pQCXIB retroviral construct for stable overexpression, using sticky ends ligation with ApaI and BamHI-HF (New England BioLabs) restriction enzymes, per manufacturer’s instructions. HEK293-FT cells were transfected with pBABE (empty vector), pBABE-G106V *PIK3CA,* pQCXIB (empty vector) and pQCXIB -S37F *CTNNB1* using Fugene 6 (Promega) per manufacturer’s instructions. Virus containing media was harvested at 24 hrs and 48 hrs post-transfection. HCC827 cells were infected with virus containing media, supplemented with 8 μg/ml of polybrene (Sigma-Aldrich), for 24 hours. The culture medium was changed to standard growth media for an over-night incubation, after which cells were incubated in antibiotic selecting medium containing puromycin 1 ug/mL (Gibco) for p-Babe construct or blasticidin 2.5 ug/mL (Gibco) for pQCXIB constructs. Antibiotic resistant cells were used in the subsequent tests.

### Cell Viability and Growth Assays

One hundred thousand of HCC827 cells, engineered with the S37F b-catenin and G106V PIK3CA constructs, and under puromycin (1 ug/mL) and blasticidin (2.5 ug/mL) selection, were seeded in 12 well plates and, after 24 hrs, treated with DMSO (control), erlotinib (50-100 nM) and rociletinib (100 nM), in 2% FBS, for three days. Cells were then air-dried for 5 minutes, fixed for 5 miutes in ParaFormAldehyde (PFA, 4% vol/vol; Santa Cruz Biothechnology) and stained in 0.05% crystal violet (g/mL; Sigma-Aldrich) solution for 30 minutes. Each well was washed twice with tap water and air-dried. Plates were scanned using ImageQuant LAS4000 (GE Healthcare Life Sciences). Each image is representative of a triplicate experiment. Cell viability was assessed using the above culture conditions, seeding two hundred of cells each well. Cell count was registered after three days of growth and assessed using Vi-CELL XR. Each test was run in triplicate. One-way ANOVA and Student's T-test with Bonferroni correction were used to assess statistical significance (GraphPad Prism). The variation between the sample sets was similar and expressed as standard deviation.

### Invasion and Migration Assays

Transwell migration and invasion assays were performed as described in Okimoto *et al.*^50^ Briefly, 8-μm-pore Matrigel coated (invasion) or non-coated (migration) Transwell inserts (BD Biosciences) were added at the top of a Transwell chamber filled with 10% FBS, RPMI media. To each insert, 2.4 × 10^4^ cells in serum-free media were added. The Transwell chambers were incubated for 20 hrs at 37 °C in the incubator. Cells that did not migrate through the pore or invade the matrigel were scraped off; the membranes were fixed in methanol for 15 min and then stained with crystal violet for 30 min. The surface of the membrane was imaged in 5 distinct fields, with a Zeiss Axioplan II immunofluorescent microscope at 10×. Invasion and migration were assessed counting the average imaged cells in the 5 regions. Results represent the average of three independent tests. One-way ANOVA and Student's T-test with Bonferroni correction were used to assess statistical significance (GraphPad Prism). The variation between the sample sets was similar and expressed as standard deviation.

### Immunoblotting and q-RT-PCR

The HCC827 cells engineered with the S37F b-catenin and G106V PIK3CA constructs were drug treated, in serum free condition, with DMSO (control), erlotinib (100 nM) and rociletinib (100 nM) for 24 hrs. Protein lysates were collected in RIPA buffer supplemented with protease (Roche) and phosphatase (Roche) inhibitors. Western blot was performed loading 10 ug of lysed proteins. Pre-casted 4-15% gels (Bio-Rad) were used for the mono-dimension protein separation. Proteins were transferred on nitrocellulose membranes using Trans-blot Turbo Transfer system (Bio-Rad). Blots were then blocked in Tris-buffered saline, 0.1% Tween20 (vol/vol) and 5% BSA (Fischer Scientific, vol/vol) for 1 hr, at room temperature. The primary antibodies were incubated over-night, at 4°C. The primary antibodies used were: pY1068-EGFR D7A5 (#3777), total EGFR D38B1 (#4267), β-Catenin D10A8 (#8480), pS473-AKT D9E (#4060), total AKT (#9272), pT202/Y204-ERK1/2 (#9101), total ERK1/2 (#9102) and cleaved PARP (#9541) from Cell Signaling Technology; Actin AC-74 (#A2228) from Sigma-Aldrich. The membranes were washed twice in washing buffer (Tris-buffered saline, 0.1% Tween20, vol/vol) and then incubated with secondary HRP conjugated antibodies for 1 hr, at room temperature. ECL kit (GE Healthcare) was used as chemoluminescent substrate. Blots were developed and scanned using ImageQuant LAS4000 (GE Healthcare Life Sciences). ImageJ (NIH) was used to quantify the western blots. All western blots represent the result of three independent experiments.

The RNA was purified from the HCC827 cells engineered with the S37F b-catenin and G106V PIK3CA constructs using RNeasy Micro Kit (Qiagen). One microgram of total RNA was used for the reverse-transcriptase reaction with SensiFAST cDNA Synthesis Kit (BIOLINE). The q-PCR was performed with six replicates each condition and using a 1:3 dilution of the template cDNA. Human *c-myc, cyclin-D1, LEF-1, HOXB9,* and endogenous control *GAPDH* genes were amplified with Taqman gene expression assay (Applied Biosystems). Gene expression analysis was computed using QuantStudio 12K Flex Software (Applied Biosystems). Data were analyzed using the 2^−ΔΔ*C*t^ method and expressed as relative mRNA expression. One-way ANOVA and Student's T-test with Bonferroni correction were used to assess statistical significance (GraphPad Prism). The variation between the sample sets was similar and expressed as standard deviation. Immunohistochemistry: Immunohistochemistry was performed as described^32^. Briefly, 5-micron thick formalin-fixed paraffin embedded (FFPE) human tissue sections were stained with the β-Catenin D10A8 (#8480 Cell Signaling, 1:100 dilution), or pSer473-Akt D9E (#4060, Cell Signaling, 1:100 dilution) antibody per manufacturer’s instructions. Stained slides were digitized using the Aperio ScanScope CS Slide Scanner (Aperio Technologies) with a 20x objective. The proportion of cells exhibiting nuclear β-Catenin staining was determined using the ScanScope default nuclear algorithm. pSer473-Akt quantitation was determined using the ScanScope default membrane algorithm. Three fields of view per section were used to determine the mean and standard error of the mean of positively staining cells.

## Acknowledgments

The authors acknowledge funding support from NIH: NCI-R01CA169338, NIH Director’s New Innovator Award NCI-DP2CA174497, the Pew Charitable Trust, Stewart Foundation, and Searle Foundation (to T.G.B), and to AACR and Lung Cancer Research Foundation (C.M.B.). The authors thank Joel Blakely for artwork and Amit Sabnis, Ross Okimoto, and Asmin Tulpule for critical review of the manuscript.

## Author Contributions

C.M.B., T.B.K.W, C.S. and T.G.B designed the study. C.M.B. performed medical record review, analyzed data and prepared tables and figures. T.B.K.W. performed whole-exome sequencing and clonality analysis and prepared tables and figures with assistance from N.M., G.A.W, and N.J.B. W.W. performed analysis of cell-free DNA sequencing data on patient cohorts and prepared tables and figures. B.G. performed cell line experiments and prepared figures with assistance from A.M. J.J.C. and M.D. performed CAPP-Seq analysis. V.O. and J.R. performed IHC analysis. C.E.M, M.A.G, V.W., P.J.M, D.R.G, H.H, R.C.B, J.W.R., performed medical record review and provided clinical data. K.C.B. and R.B.L. compiled and annotated cfDNA data from 1150 EGFR-mutant positive and 1008 EGFR mutant negative NSCLC patients. A.R.C. extracted DNA and prepared exome libraries from patient tumor samples. A.F.C. and J.S.J. performed exome sequencing alignment and quality analysis. P.G. harvested autopsy tissue and performed pathological assessments. C.M.B. and T.G.B. wrote the manuscript with contributions from all authors.

## Author Information

Reprints and permissions information is available at http://www.nature.com/reprints.

K.C.B. and R.B.L. are employees of Guardant Health Inc., A.F.C., J.S.J., A.R.C. and P.G. are employees of Driver Inc. Correspondence and requests for materials should be addressed to Trever G. Bivona MD PhD or Charles Swanton MD PhD.

